# Assessing the Role of Marker Density and Minor Allele Frequency on Machine Learning–Driven Genomic Selection Accuracy in Grapevine

**DOI:** 10.64898/2026.07.11.737951

**Authors:** Felipe Roberto Francisco, Geovani Luciano de Oliveira, Guilherme Francio Niederauer, Roberto Fritsche-Neto, Anete Pereira de Souza, Mara Fernandes Moura Furlan

**Author notes:** Corresponding author, (MFMF).

## Abstract

Although grapevine (*Vitis* spp.) is among the oldest and most economically significant fruit species globally, its genetic improvement faces major bottlenecks due to long juvenile periods and extended cycles for phenotypic evaluation. In this context, genomic selection (GS) has emerged as an effective alternative to traditional selection, offering a robust framework to optimize breeding programs by significantly reducing generation intervals while enhancing predictive accuracy (PA) in early generations and expected genetic gains (EGGs). Nevertheless, factors such as minor allele frequency (MAF) and population size can significantly affect predictive models, even to the point of making their use unfeasible in breeding programs. In this context, this study evaluated the effect of data dimensionality reduction on GS accuracy by selecting single-nucleotide polymorphisms (SNPs) based on MAF thresholds. The experimental design tested the predictive capacities of four machine learning (ML) algorithms (ElasticNet, K-Neighbors, Support Vector Machine Regression, and XGBoost) alongside the conventional Genomic Best Linear Unbiased Prediction (gBLUP) model. These were validated using three SNP datasets (11,115, 9,494, and 6,100 markers) filtered by MAF levels of 0.05, 0.1, and 0.2 across six genetic traits, and EGGs were compared between conventional breeding and GS via the breeders’ equation. The results revealed that the ML models exhibited remarkable stability, with no significant differences in PA across the different MAF-based SNP densities, except for berry length, which showed a substantial difference with XGBoost at an MAF of 0.2. Conversely, gBLUP demonstrated high sensitivity to dimensionality reduction, with its performance significantly impacted by MAF filtering across all the traits. These results suggest that compared with traditional GS models that rely on a genomic kinship matrix, ML-based approaches offer greater flexibility in feature reduction. Additionally, compared with chemical traits, morphological traits generally had greater predictive ability. Furthermore, every GS model provided estimated genetic gains superior to traditional breeding, with improvements ranging from an 8.90-fold increase in berry length to a 2.86-fold increase in total soluble solids, confirming that GS integration is promising for enhancing breeding efficiency in grapevines.

## Introduction

Grapevine (*Vitis* spp.) is one of the oldest and most significant fruit species worldwide, with cultivation dating back to the Bronze Age, approximately 3,500 BCE (Leão & Possídio, 2000; Jiang et al., 2009; Maras et al., 2020; Candar et al., 2021). Although viticulture is an exotic species in Brazil, it has become increasingly vital to national agriculture, with a cultivated area of approximately 84,380 ha and an average yield of 21,570 kg ha^-1^ (IBGE, 2024). The success of Brazilian viticulture is largely attributed to genetic improvement, which has enabled the expansion of grapevine cultivation from temperate regions into tropical zones (Camargo et al., 2011). However, national viticulture still faces significant challenges, particularly with respect to the subtropical climate characterized by mild, cold, dry winters and hot, humid summers. These conditions can hinder the dormancy phase due to insufficient chilling, leading to hormonal imbalances that may adversely affect fruit quality and yield (Moura 2010; Mariani et al., 2019).

Despite the significant advances in grapevine breeding, these programs still face challenges, primarily due to the long breeding cycle (Alleweldt & Possingham, 1988; Camargo et al., 2011; Bharati et al., 2023). However, advances in molecular biology have enabled marker-assisted selection (MAS), which uses molecular markers to select individuals early within a population (Song et al., 2023). This strategy has been adopted by several breeding programs across different species (Beyene et al., 2021; Calleja et al., 2022; Lu et al., 2025)

Linkage mapping (Qi et al., 2021; Miao et al., 2022; Jiang et al., 2024) and genome-wide association studies (GWASs, Francisco et al., 2019; Sahito et al., 2024; Wang et al., 2024; Xue & Cui, 2025) are commonly used to identify genes or quantitative trait loci (QTLs) associated with traits of interest, providing molecular markers that can subsequently be applied in MAS. Although this QTL identification strategy is widely used across different species and breeding programs, it presents limitations regarding the quantitative nature of many yield traits, which are generally complex and associated with many genes; therefore, they individually have a small effect on the phenotype, hindering their use in breeding programs (Francisco et al., 2019). For these complex traits, genomic selection (GS) is an interesting alternative, as it uses a large number of markers without prior knowledge of their effect on the phenotype to predict the performance of individuals in a population, enabling the early selection of superior individuals (Souza et al., 2019; Francisco et al., 2021; Aono et al., 2022; Bharati et al., 2023). Traditionally, GS methods are based on frequentist or Bayesian models, which require strong assumptions or a priori information, such as the dominant, additive, or epistatic nature of the traits, which are often difficult to model, making them less common in practice (Tibshirani, 1996; Meuwissen et al., 2001; VanRaden, 2008; Pérez & Los Campos, 2014; Pérez-Enciso & Zigaretti, 2019). Additionally, they are unable to capture the nonlinear effects expected when machine learning (ML) algorithms are used (Shokor et al., 2025).

A second aspect that remains under discussion is data preprocessing, particularly given the high dimensionality introduced by the vast number of markers enabled by advances in next-generation sequencing (NGS) technologies. These technologies allow the identification of thousands or millions of single-nucleotide polymorphism (SNP) markers, in contrast to the small number of observations (i.e., phenotyped individuals), an effect known as ‘large *p*, small *n*’ (Long et al., 2007; Konietschke et al., 2021; Dhal et al., 2022). As a proposal to circumvent such effects, feature selection techniques can be used to select the best predictors from the dataset, potentially yielding significant gains in prediction performance and increasing data processing efficiency (Konietschke et al., 2021; Dhal et al., 2022). A method still underexplored in the literature for selecting the best predictors for GS may be filtering by minor allele frequency (MAF), as low-frequency alleles in the training population can bias model inference (Uemoto et al., 2015; Linck & Battey, 2019). Furthermore, markers in high-density panels often exhibit strong linkage disequilibrium, resulting in high multicollinearity (Gianola et al., 2003; Al-Mamun et al., 2024). This redundancy makes the models sensitive to noise and considerably increases the risk of overfitting (Pérez-Enciso, & Zingaretti, 2019, Al-Mamun et al., 2024).

The combination of factors such as the inclusion of markers with small MAFs, insufficient population size, and trait distribution negatively influences predictive accuracy, mainly due to the biased estimation of SNP effects (Zhang et al., 2019). Furthermore, these low-frequency markers may result from sequencing artifacts and are often reported to lack significant contributions to the genetic variation of the population (Purcel et al., 2007; Lee et al., 2024). Therefore, all these characteristics make markers with small MAFs good candidates for removal, thereby reducing data dimensionality and increasing computational efficiency without compromising predictions.

In this context, beyond dimensionality reduction via feature selection, another significant challenge concerns the selection and tuning of ML models, as different approaches model the relationships between genotypes and phenotypes in distinct ways (Chen et al., 2004). This stage is fundamental for ensuring high predictive accuracy (PA) and becomes even more critical given the complexity of genomic data and the diversity of available modeling architectures.

Wide ranges of ML models and parameters are available that can significantly improve genomic prediction (GP), enabling early and accurate selection in breeding programs (Liang et al., 2022). However, choosing the most appropriate model and tuning its parameters to the dataset represents a complex task, given the number of elements to be considered, such as the number of hidden layers, neurons per layer, learning rate, type and number of filters, activation functions, optimization algorithms, and regularization strategies (Montesinos et al., 2021; Galli et al., 2022). The complexity of these choices is one of the main obstacles to the use and popularization of these tools, as implementing these technologies in breeding programs is challenging, mainly due to the need for in-depth technical knowledge in modeling, automation, and programming languages (Waring et al., 2020; Montesinos et al., 2021). To overcome these barriers, several tools, such as AutoWEKA (Thornton et al., 2013), hyperopt-sklearn (Komer et al., 2019), Auto-sklearn (Feurer et al., 2022), TPOT (Olson et al., 2016), and Auto-Keras (Jin et al., 2019), have been developed based on the automated machine learning approach, which performs the automatic selection of models and parameters most suitable for the dataset (Mahima et al., 2021; Galli et al., 2022). Nevertheless, these solutions require significant computational power and can take a long time, as they perform exhaustive searches over different combinations of hyperparameters to find the best-performing configuration for the task at hand.

Some ML methods are more effective for different tasks, such as support vector machines (SVMs) (Vapnik et al., 1995) and random forests (Breiman et al., 2001) for classification. Nevertheless, these methods must also have their parameters adjusted to the problem specifics. This fine-tuning task can be simplified using the (grid search) method (Bergstra et al., 2012), which combines different parameters presented by the researcher until the best model is obtained within the combinations of this set. This may be the most efficient method for identifying the best model, as it integrates the researcher’s prior knowledge of the data with automated grid search. In plant breeding, these ML tools, whether automated or not, have shown promising results across different crops, such as maize, soybean, and rubber tree (Aono et al., 2022; Galli et al., 2022; Sarkar et al., 2023). Therefore, the use of these tools for grapevine breeding can provide great benefits.

In this context, this work employs various ML strategies and traditional GS methods to optimize and improve the efficiency of grapevine breeding programs. Here, we tested different unsupervised ML models to identify genetic groups based on six agronomically important grapevine traits. Additionally, we trained ML algorithms and traditional genomic selection methods to reduce marker density and bias by optimizing molecular marker filtering based on MAF. The results presented herein provide significant contributions by elucidating how the MAF influences different predictive models, thereby contributing to the discussion regarding the use of low-density sequencing to reduce costs in breeding programs.

## Materials and Methods

### Plant Material

The population, composed of accessions from the Active Germplasm Bank (BAG) of the Instituto Agronômico de Campinas (IAC), is located at the Center for Advanced Fruit Research and Development - IAC in Jundiaí, São Paulo, Brazil. The BAG was previously characterized for its genetic diversity (de Oliveira et al., 2020, 2023). From the total germplasm (420 genotypes), 288 distinct genotypes were selected for the GP analyses. Therefore, the chosen population consists of *V. vinifera* varieties, interspecific hybrids, and other *Vitis* spp. species. Each genotype consists of 3 or 4 vegetatively propagated plants grafted onto the ‘IAC 766 Campinas’ rootstock. The plants were trained in a vertical shoot positioning system, with annual pruning performed in July and maintaining only two buds per branch.

### Phenotyping

All plants were evaluated over 12 productive seasons for berry length (BL, cm), cluster length (CL, cm), cluster width (CW, cm), rachis fresh mass (RFM, g), total soluble solids (TSS, °Brix), and the maturation index (MI), calculated as the ratio between total soluble solids and titratable acidity. All these phenotypes exhibited Pearson correlations of less than 0.7.

The trait means per season were adjusted using mixed models to obtain the best linear unbiased estimates (BLUEs) from equation 1, implemented in the sommer package in R (Covarrubias-Pazaran, 2018). These adjusted means were then utilized for the GP analyses.

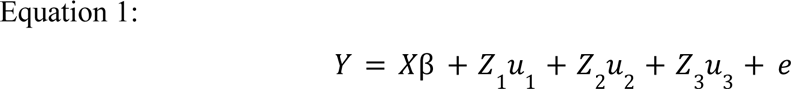

where *Y* represents the observed values in the field; *X*β denotes the fixed effects of the model, in which β includes the genotype effects; *Z*_1_ *u*_1_ represents the random effects of genotypes considering the evaluation field matrix; *Z*_2_ *u*_2_ corresponds to the random effects of rows; and *Z*_3_ *u*_3_ represents the random effect structure for columns across the three experimental fields. Finally, *e* denotes the residuals, which are assumed to be *e* = *N*(0, *I*δ^2^).

The best linear unbiased predictions (BLUPs) (Equation 2) were estimated using the sommer package in the R environment (Covarrubias-Pazaran, 2018) to estimate broad-sense heritability (*H*^2^), as demonstrated in Equation 3.

The following equation can describe the statistical model used:

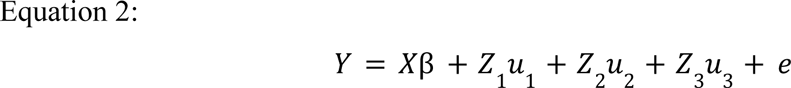

where *Y* represents the response variable, the values measured in the field; *X*β corresponds to the fixed effect of the field; *Z*_1_ *u*_1_ represents the random effects of genotypes within each field with field-specific variance; *Z*_2_ *u*_2_ represents the random effects of rows within the fields; *Z*_3_ *u*_3_ represents the random effects of columns, considering only fields “A” and “C”; and *e* is the vector of random errors, assumed as *e* = *N*(0, *I*δ^2^).

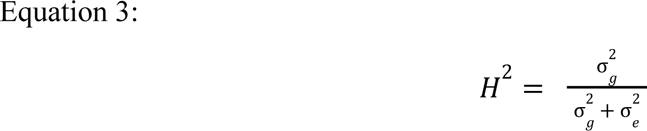

where *H*^2^ represents the broad-sense heritability; σ^2^*_g_* is the genetic variance (sum of genotypic effects within the environment); and σ^2^*_e_* is the residual variance (experimental error).

### Identification of Contrasting Phenotypic Groups

To identify the best individuals in the population based on phenotypic sets (groups), we performed phenotype clustering. To allow a fair comparison of these traits, normalization was applied using the min–max method via the rescale function in R.

The optimal number of groups was determined from the data using the NbClust algorithm (Charrad et al., 2014), with k ranging from 2–30. The evaluation used the Euclidean distance and the Silhouette criterion to determine the optimal number of clusters. Based on these criteria, k=3 was identified as the ideal number of clusters, which was subsequently applied in the clustering stage using the k-means algorithm (MacQueen, 1967).

### DNA Extraction and Genotyping

Total genomic DNA was extracted from leaf tissue using a DNeasy® Plant Mini Kit (QIAGEN, Hilden, Germany) according to the manufacturer’s instructions. DNA quality was assessed using 1% agarose gel electrophoresis and a NanoDrop 8000 spectrophotometer (Thermo Fisher Scientific, Wilmington, DE, USA). The DNA concentration was subsequently determined using a Qubit 3.0 fluorometer (Thermo Fisher Scientific, Waltham, MA, USA).

The 288 selected accessions were genotyped using the commercial Vitis18kSNP array at Neogen (Pindamonhangaba, Brazil), following the Infinium HD Assay Ultra protocol (Illumina Inc., San Diego, CA, USA). Raw data were visualized and clustered using GenomeStudio 2.0 software (Illumina Inc., San Diego, CA, USA). The SNP filtering tool ASSIST 1.02 (Di Guardo et al., 2015) was subsequently used, employing default thresholds adapted for germplasm material to filter the dataset. SNPs classified as ‘Monomorphic’, ‘Failed’, and ‘NullAllele-Failed’ were removed (Di Guardo et al., 2015).

The markers were subsequently filtered using snpReady software (Granato and Fritsche-Neto, 2018). For dimensionality reduction, three distinct minor allele frequencies (MAFs) of 0.05, 0.10, and 0.20 were used; this strategy was adopted to reduce potential population bias caused by allele sampling in small populations (Linck & Battey, 2019). Furthermore, using snpReady software (Granato and Fritsche-Neto, 2018), the markers were filtered with a minimum call rate of 0.85, individuals with more than 0.99 missing SNPs were removed, and missing data were imputed using the k-nearest neighbors (kNN) algorithm (Hastie et al., 2020). We used the native R function (prcomp) for principal component analysis (PCA) calculation, and a graphical representation of the results was created using the ggplot2 package (Wickham, 2016) to identify potential data structures that could compromise the accurate understanding and interpretation of the GP results (Aono et al., 2022).

### Training of Predictive Models

To train the predictive model, the dataset was partitioned into training (90%) and test (10%) sets using the train_test_split function from the scikit-learn library (Pedregosa et al., 2011). Using only the training set, we trained the GP models using the conventional genomic selection method, Genomic Best Linear Unbiased Prediction (gBLUP), which is based on the VanRaden additive relationship matrix (VanRaden, 2008) created with snpReady software (Granato and Fritsche-Neto, 2018). In addition to this method, we utilized the machine learning models ElasticNet, K-Nearest Neighbors, Support Vector Regression (SVR), and XGBoost, which are implemented in the scikit-learn library (Pedregosa et al., 2011) in Python. These machine learning models were selected for their nonparametric nature, which provides high flexibility for identifying complex associations between molecular markers and the phenotypes of interest (Montesinos-López et al., 2021).

ElasticNet was chosen because it is a linear regression model with L1 and L2-norm regularization of the coefficients and is primarily used when many correlated attributes are observed in the dataset (Koh et al., 2007; Friedman et al., 2010). K-Nearest Neighbors, modified for regression, was implemented as a distance-based method, a simple and flexible approach capable of handling different data types and adapting to irregular features (Yao & Ruzzo, 2006). SVR is defined as a supervised machine learning model with high GP potential (Mota et al., 2024). Finally, XGBoost (Chen & Guestrin, 2016) is a highly promising and effective decision tree-based method for genomic prediction (Elgart et al., 2022).

The best hyperparameters were determined using the Grid Search method (Bergstra, 2012), which exhaustively explores a predefined set of parameters (Supplementary Table 2). Cross-validation was performed using only the best hyperparameters identified by Grid Search, employing a stratified k-fold method repeated 50 times, and the mean Pearson correlation between observed and predicted BLUEs was used as the model’s predictive accuracy (PA). Differences between the models and datasets, based on the MAF, were tested using permutation tests (Ojara and Garriga, 2010) via the R package lmPerm (Wheeler & Torchiano, 2025). Sets with significant differences were tested pairwise using the coin package (Hothorn et al., 2006) based on adjusted p values. Finally, all the models were evaluated on the final test set; this set was never used for hyperparameter tuning or during model training and thus served as an external dataset, allowing for the assessment of model consistency on new data.

### Expected Genetic Gains

To compare the expected genetic gains based on traditional genotype selection methods (*EGG_c_*) and expected genetic gains using GP models (*EGG_pa_*), we used the breeder’s equation (Grattapaglia 2017; Matias et al., 2019; Souza et al., 2019), which simulates genetic improvement in a traditional selection scenario (Equation 4) and is also based on the use of markers and GP models (Equation 5).

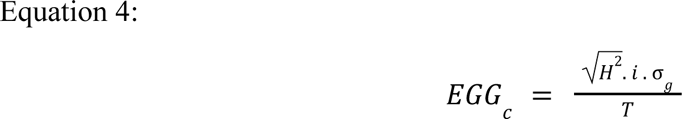

where *H*^2^ is the broad-sense heritability; *i* is the selection intensity; σ*_g_* is the additive genetic standard deviation; and *T* is the selection cycle time (15 years in traditional breeding).

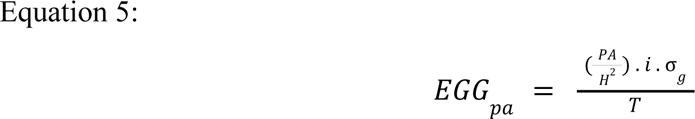

where *EGG_pa_* is the expected genetic gain per selection cycle; PA is the Pearson accuracy of the models; i is the selection intensity; H² is the broad-sense heritability; σ*_g_* is the additive genetic standard deviation; and T represents the time for each selection cycle considered (2 years).

For classical breeding, we estimated a selection cycle of 15 years, which is the average time for genotype selection in grapevine breeding programs (Eibach & Töpfer, 2015), and 2 years when using GS tools, the time required to obtain genetic material without causing the individual’s death.

## Results

### Genotyping and Phenotyping

A total of 12,264 SNPs were identified in the population. After quality control filtering, three new datasets were constructed based on the MAF thresholds (Table 1). The number of markers decreased substantially as more restrictive MAF thresholds were applied, ranging from 11,115 markers for an MAF > 0.05 to 6,100 markers for an MAF > 0.20, a reduction of approximately 55% of the SNPs. Imputation rates remained low, with a maximum of 1.82%, and removing low-quality markers did not improve them, indicating that the final dataset was high-quality and consistent for subsequent analyses. Based on this dataset (11,115 SNPs), no clear population structure was observed, as indicated by the sum of the first two principal components, which explained 17.3% of the total variation (Supplementary Figure 2).

**Table 1:**
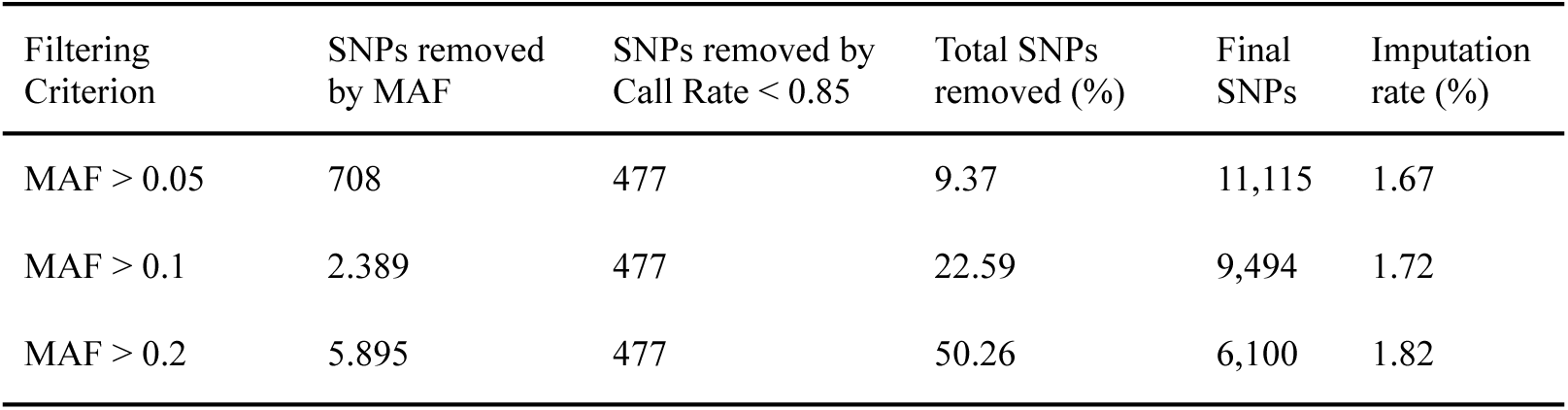
Final marker sets obtained for the different datasets based on MAF thresholds.

| Filtering Criterion | SNPs removed by MAF | SNPs removed by Call Rate < 0.85 | Total SNPs removed (%) | Final SNPs | Imputation rate (%) |
| --- | --- | --- | --- | --- | --- |
| $MAF > 0.05$ | 708 | 477 | 9.37 | 11,115 | 1.67 |
| $MAF > 0.1$ | 2,389 | 477 | 22.59 | 9,494 | 1.72 |
| $MAF > 0.2$ | 5,895 | 477 | 50.26 | 6,100 | 1.82 |

The k-means algorithm identified three contrasting phenotypic groups (Figure 1). The first two principal components, PC1 (39.4%) and PC2 (25.9%), together explained 65.3% of the total variation present in the data. Furthermore, apparent clustering, which was related mainly to the nature of the evaluated traits (chemical or morphological), was evident (Figure 1). An overlap of individuals at the centroid, which may indicate difficulty in discriminating between these genotypes, could also be observed (Figure 1).

**Figure 1:**
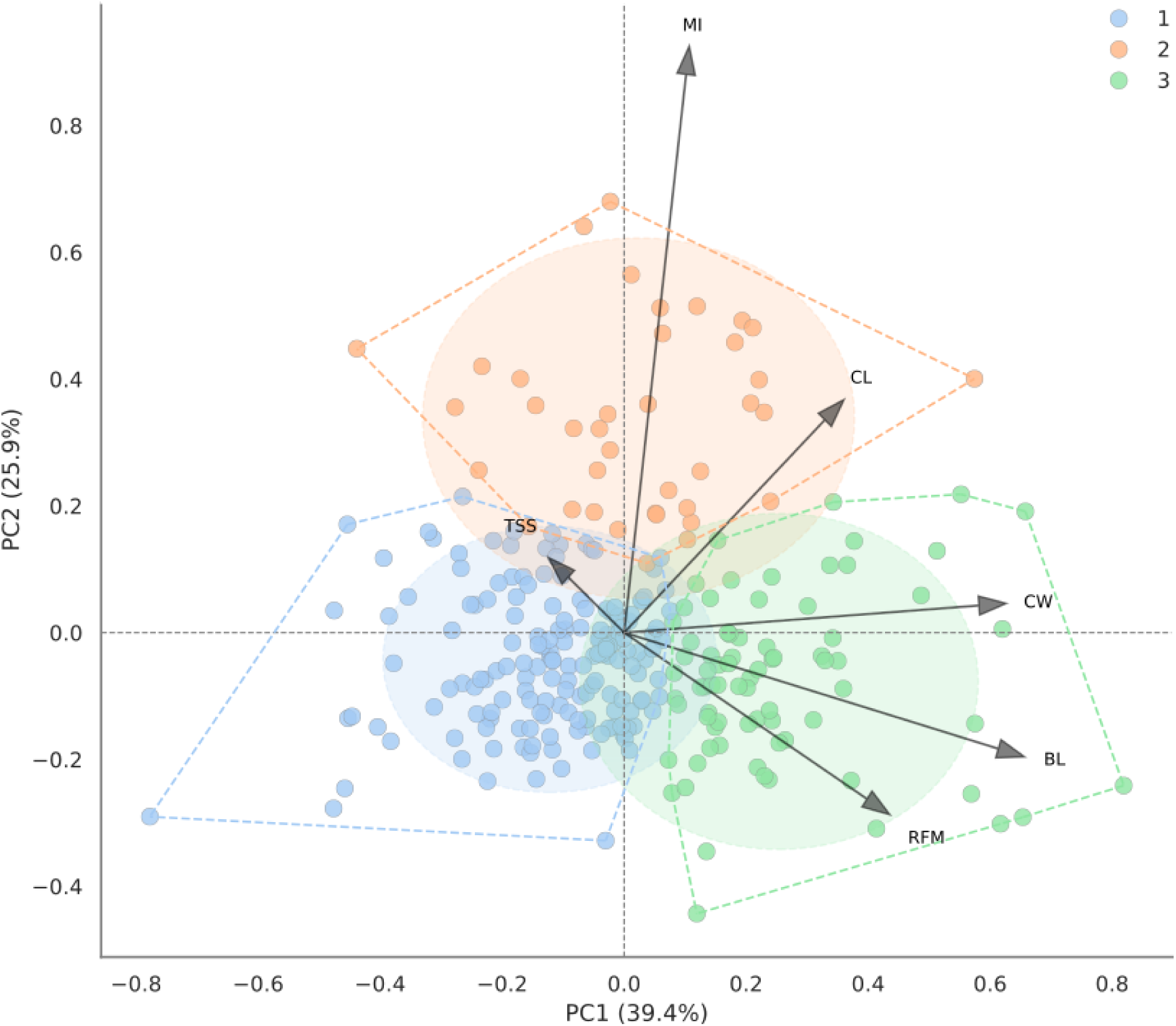
First principal component analysis (PCA) based on adjusted phenotypes, showing the three contrasting phenotypic groups identified using the k-means algorithm. The characteristics are abbreviated as berry length (BL, cm), cluster length (CL, cm), cluster width (CW, cm), rachis fresh mass (RFM, g), total soluble solids (TSS, °Brix), and maturation index (MI).

The analyzed traits exhibited moderate to high H² values, indicating a significant genetic contribution to the observed phenotypic variation. The estimated H² values were 0.57 for CL and MI, 0.70 for TSS, 0.77 for CW, 0.82 for RFM, and 0.94 for BL.

### Prediction Models

No substantial differences in mean predictive accuracy were observed among the evaluated GP models, although substantial differences were detected for specific traits and MAFs. In contrast, apparent differences in PA were observed, primarily related to the nature of the evaluated traits. Compared with chemical characteristics, morphological traits generally exhibited higher mean predictive accuracies (Figure 2).

**Figure 2.**
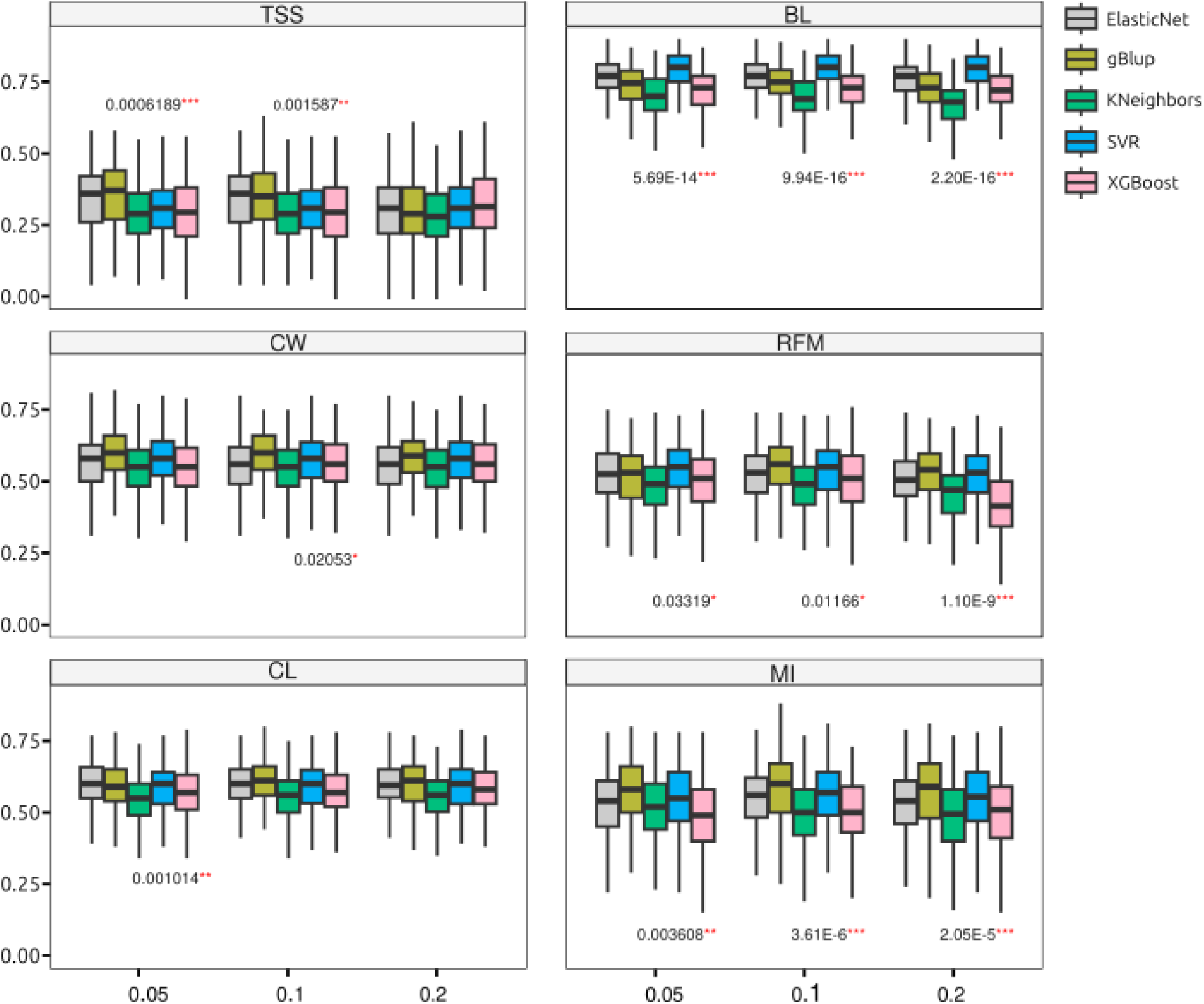
Comparison of predictive accuracies among the tested models. Overall adjusted p values indicate significant differences between models within the same MAF level (x-axis). Abbreviations: berry length (BL, cm), cluster length (CL, cm), cluster width (CW, cm), rachis fresh mass (RFM, g), total soluble solids (TSS, °Brix), and maturation index (MI).

Among the morphological traits, BL had the highest accuracy, ranging from 0.79 (SVR – MAF 0.05) to 0.67 (K-Nearest Neighbors – MAF 0.20). This trait showed significant differences in PA across the MAFs and models, as assessed by the permutation test (Supplementary Table 4 and Figure 2). This was followed by CW, whose accuracies oscillated between 0.61 (gBLUP – MAF 0.10) and 0.54 (K-Nearest Neighbors – MAF 0.05) and showed significant differences in PA among the models, but only for the dataset with an MAF of 0.10 (Supplementary Table 4 and Figure 2). CL presented PA values between 0.60 (gBLUP – MAF 0.05) and 0.54 (K-Nearest Neighbors – MAF 0.10 and 0.20; XGBoost – MAF 0.05), with significant differences observed only for the MAF 0.05 dataset. Finally, the RFM trait exhibited accuracies ranging from 0.55 (gBLUP – MAF 0.10) to 0.41 (XGBoost – MAF 0.20), and the predictive accuracies significantly differed across the other datasets, although these differences were small and nonsignificant in the pairwise test, with adjusted p values for MAFs of 0.05 and 0.10 (Supplementary Table 4 and Figure 2). RFM was also the only morphological trait whose performance was lower than that of MI, a chemical trait.

Among the chemical traits, MI had predictive accuracies ranging from 0.58 (gBLUP – MAF 0.10) to 0.48 (K-Nearest Neighbors – MAF 0.20), with significant differences in predictive accuracy across the models and datasets (Supplementary Table 4 and Figure 2). Finally, the trait with the lowest predictive performance was TSS, with values ranging from 0.35 (gBLUP – MAF 0.05) to 0.28 (K-Nearest Neighbors – MAF 0.20). Major differences in predictive accuracy across datasets and models were detected (Supplementary Table 4 and Figure 2).

When significant differences were compared within the same model across the different MAFs (0.05, 0.10, and 0.20), only gBLUP showed significant differences across all the traits, with overall lower accuracy in the dataset with an MAF of 0.20 (Supplementary Table 5). In addition to gBLUP, the XGBoost model for the RFM trait also showed significantly lower differences in the dataset, with an MAF of 0.20 (Supplementary Table 5).

With respect to the test set, the TSS showed the greatest difference in predictive accuracy between the validation and test sets; this difference was 0.43 for the TSS–gBLUP–MAF 0.20 model and 0 for the TSS–XGBoost–MAF 0.05 model. The CL trait also showed zero difference between the two sets (CL – gBLUP – MAF 0.10). In general, the discrepancies observed between training and test performance were low, highlighting the consistency and predictive stability of the evaluated models (Supplementary Table 6).

### Expected Genetic Gains

Compared with those obtained through classical breeding, all the evaluated genomic selection models presented higher EGGs (Figure 3). With respect to TSS, which was the trait with the lowest predictive accuracy, genomic selection resulted in EGGs ranging from 2.86 (K-Nearest Neighbors – MAF 0.20) to 3.58 times (gBLUP – MAF 0.05) those of the classical method. For CL, these values oscillated between 5.92 (K-Nearest Neighbors – MAF 0.10 and 0.20) and 6.58 (gBLUP – MAF 0.05). With respect to MI, the improvements ranged from 6.32 (K-Nearest Neighbors – MAF 0.20) to 7.63 (gBLUP – MAF 0.10). The RFM trait showed gains ranging from 4.06 (XGBoost – MAF 0.20) to 5.44 (gBLUP – MAF 0.10). With respect to CW, the EGGs obtained via genomic selection outperformed those obtained from classical breeding, ranging from 4.76 (K-Nearest Neighbors – MAF 0.05) to 5.38 (gBLUP – MAF 0.10). Finally, the model that presented the greatest expected genetic gain relative to classical breeding was BL, with values ranging from 6.86 (K-Nearest Neighbors – MAF 0.20) to 8.90 (SVR – MAF 0.05, 0.10, and 0.20).

**Figure 3:**
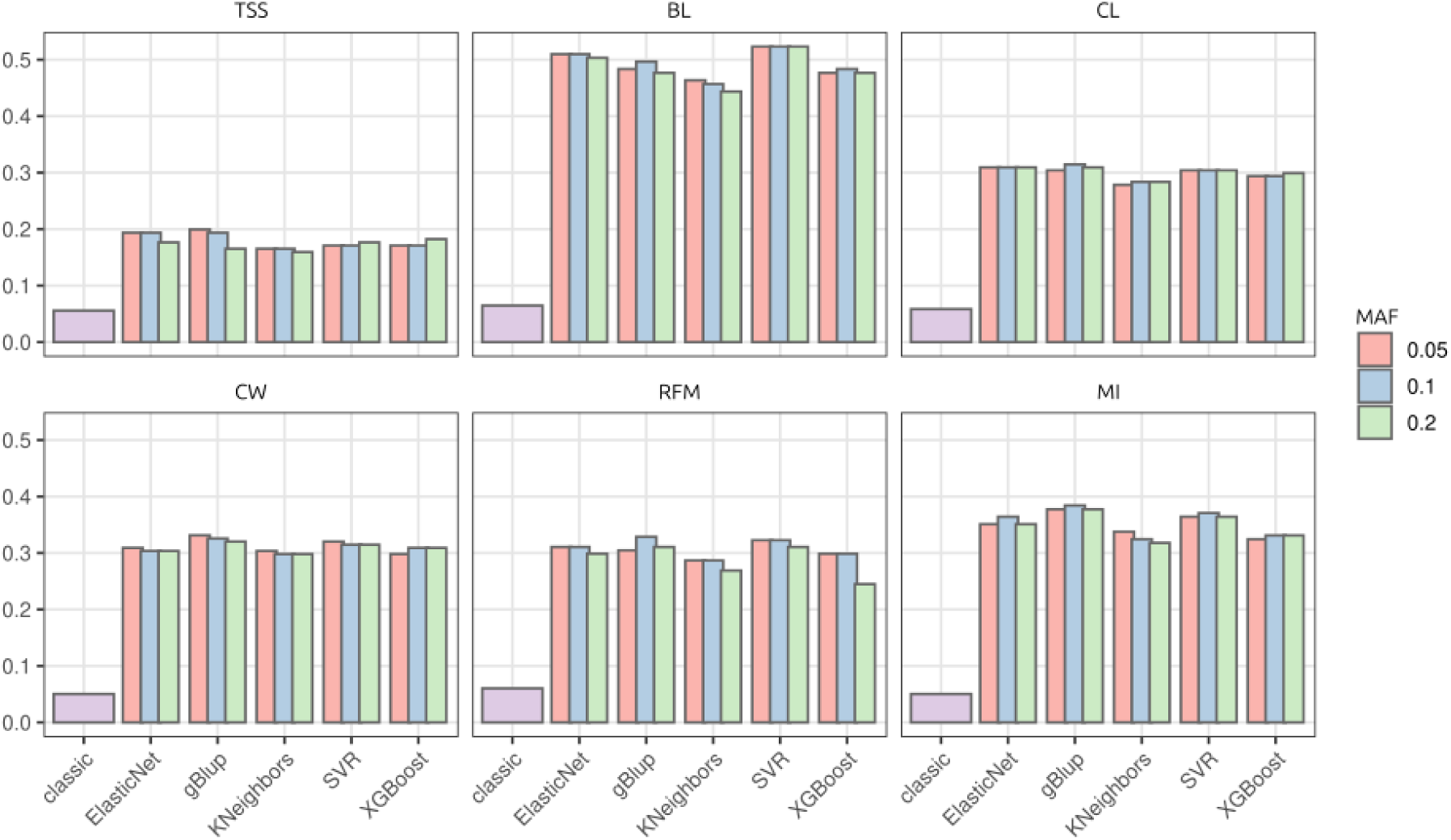
Expected genetic gains (EGGs) based on the use of genomic selection tools and classical breeding. For each model, EGGs are shown for datasets with minor allele frequencies (MAFs) of 0.05, 0.10, and 0.20. Abbreviations: berry length (BL, cm), cluster length (CL, cm), cluster width (CW, cm), rachis fresh mass (RFM, g), total soluble solids (TSS, °Brix), and maturation index (MI).

## Discussion

### Phenotyping

Traditionally, the genetic improvement of perennial species, such as grapevine (*Vitis* spp.), is based on phenotypic evaluation. The success of this selection depends intrinsically on the trait’s heritability, which decomposes the observed phenotypic variation into genetic and environmental components, as well as the genotype-by-environment interaction (Schmidt et al., 2019). In breeding, heritability quantifies the proportion of total phenotypic variance attributable to genetic variation, thereby determining the efficiency with which an individual’s genetic potential is transmitted to subsequent generations (Schmidt et al., 2019).

Heritability was high for all the traits, exceeding 0.57, indicating strong selection potential (Amegbor et al., 2022). We also highlighted that unsupervised ML methods for identifying contrasting phenotypic groups within a germplasm have proven promising (Dallastra et al., 2014; Oliveira et al., 2016). This strategy can significantly increase the efficiency of plant breeding programs by facilitating the selection of contrasting parents and maximizing genetic variability in the evaluated populations (Dallastra et al., 2014; Oliveira et al., 2016). The use of the NbClust algorithm (Charrad et al., 2014) to identify the optimal number of phenotypic clusters enabled the automation of this process while reducing potential researcher bias (Charrad et al., 2014). Based on this strategy, three distinct phenotypic groups were identified using the k-means algorithm (MacQueen, 1967). These clusters exhibited contrasting characteristics, suggesting that they could be applied in plant breeding programs, especially for selecting divergent parents. The results obtained corroborate those of Oliveira et al. (2020), who genetically characterized this germplasm using microsatellite markers and classical population inference methods, such as STRUCTURE (Pritchard et al., 2000).

Such results can positively impact breeding programs for the species, as most of these programs aim to cross divergent genotypes to obtain hybrids that combine the best traits of their parents. Here, we present an approach that uses simple unsupervised ML tools to identify contrasting phenotypic groups within a diverse grapevine germplasm.

### Genomic Selection in Grapevine

Grapevine breeding requires an extended time frame, potentially taking up to 30 years from a controlled cross to the release of an improved genotype (Eibach et al., 2015; Mota et al., 2024). This is primarily due to the time required for field evaluations (Eibach et al., 2015). In this context, GS becomes a powerful tool, as it can be implemented as soon as molecular markers are obtained without causing the death of the individual (Grattapaglia et al., 2013). Several authors have demonstrated that such a tool is particularly relevant for the breeding of perennial species, such as in the case of grapevines (Grattapaglia et al., 2013; Souza et al., 2019; Aono et al., 2022; Brault et al., 2023). For genomic selection to be viable, the PA must be high, which depends on the genetic nature of the trait being analyzed, the data quality, and the model used (Mota et al., 2024).

Different predictive models can capture genetic effects in distinct ways. Traditional genomic selection methods use matrices such as the VanRaden and Gaussian Kernel matrices to capture additive, dominant, and epistatic effects, which can be either summed or partitioned (VanRaden, 2008; Cuevas et al., 2016; Zhang et al., 2019). In this sense, several authors argue that the use of ML models can be an interesting alternative, as they offer advantages, mainly in their flexibility across different data distributions and their strong capacity to model nonlinear relationships (Pérez-Enciso & Zigaretti, 2019; Azodi et al., 2019). Here, we compared the predictive accuracy across various GS models, ranging from traditional models such as gBLUP, based on the additive matrix of VanRaden (2008), to nonparametric ML-based models. Our results revealed significant, although slight, differences among the PAs observed across the different models used, which may be related to the genetic nature of the evaluated traits (Mota et al., 2024). We highlight the K-Nearest Neighbors model as the worst performer among the tested models; this must be related. This may be due to the difficulty of this model in consistently capturing significant marker effects for phenotype definition (Zhang et al., 2019). With respect to the models with higher predictive accuracies, we noted that gBLUP, SVR, and ElasticNet stood out for different traits, a trend also observed by other authors (Long et al., 2011, Zhang et al., 2019, Wang et al., 2019, Mota et al., 2024). gBLUP is traditionally used for genomic selection and has proven effective across various species, including rubber trees, maize, and eucalyptus (Tan et al., 2017; Souza et al., 2019; Wang et al., 2025). With respect to the ML models, SVR and ElasticNet were shown to be good predictors in grapevine; these models have also been shown to be effective in simulated and real populations of both animal and plant species (Long et al., 2011; Oguto et al., 2012; Wang et al., 2022).

A wide variation in predictive accuracy was observed across the evaluated traits and marker sets. Among the traits, TSS presented the lowest PA values; these results were also reflected in the PA obtained in the test set. In sugarcane, using ML models such as random forest and SVR, as well as classical models (Islam et al., 2022), reported similar predictive accuracies for this trait. Environmental conditions strongly influence TSS and can vary among grape varieties depending on the rootstock used, hindering the prediction of this trait (Miele et al., 2017; Islam et al., 2022). Another trait that also showed moderate PA was RFM. These results indicate that nonlinear models better capture the complexity of the data. This trait is also quite responsive to environmental effects, which explains these results (Correa et al., 2014). Despite these relatively low predictive accuracy values, when we compared the expected genetic gains obtained using GS relative to conventional breeding methods, we observed great potential in the models presented here, with EGG values 2.86 (K-Nearest Neighbors – MAF 0.20) to 3.58 (gBLUP – MAF 0.05) times higher than those of classical breeding for TSS, whereas for RFM, these values ranged from 4.06 (XGBoost – MAF 0.20) to 5.44 (gBLUP – MAF 0.10).

Although CL showed moderate PA, this trait exhibited a high EGG, indicating a significant advantage for predictive models in breeding programs over selection based on classical methods (Souza et al., 2019). Aspects related to cluster architecture are important for grapevine breeding programs, as these traits are directly involved in berry quality, disease susceptibility, and yield (Fermaud et al., 2001; Tello et al., 2018; Brighenti et al., 2020; Torres Lomas et al., 2024). MI also showed moderate predictive accuracy, similar to that of TSS, and was strongly influenced by genotype‒environment interactions and crop management (Pereyara et al., 2025), which hindered its prediction by the models presented here. Finally, the trait that showed the highest PA was BL (0.79 SVR – MAF 0.05 to 0.67 K-Nearest Neighbors – MAF 0.20). This trait is primarily related to productivity and fruit quality, ultimately affecting the quality of derived products (Holt et al., 2008).

### The Effect of Minor Allele Frequency (MAF) on Predictive Models

The discovery of molecular markers from NGS data has enabled the identification of many SNPs using techniques such as genotyping-by-sequencing (GBS) or whole-genome sequencing (WGS) (Elshire et al., 2011; Sahl et al., 2016). Despite their importance in molecular biology, the markers identified in distinct populations are often different, making it impossible to apply models trained on one dataset to a new population. In this regard, the use of commercial SNP chips, such as the one used here (Vitis18kSNP), offers advantages by providing a fixed set of SNPs that remains consistent across populations (Wu et al., 2025), thereby enabling the application of predictive models to new populations. Furthermore, MAF can introduce bias into predictive models, affecting both population structure estimates and the PA of the models (Zhang et al., 2019). However, SNP selection based on MAF is frequently used in several studies and is generally limited to a minimum MAF of 0.05 (Souza et al., 2019; Piles et al., 2021; Hu et al., 2022).

In this study, we did not observe significant differences in predictive accuracy when ML models across different MAF thresholds for most traits were compared, except for the RFM and XGBoost models, which showed a substantial reduction in predictive accuracy at an MAF of 0.20. These results may be due to the high flexibility of these ML models and their ability to handle the exclusion of features with minor effects on the predicted trait. Similar results regarding dimensionality reduction based on different strategies have been observed in systems such as heart disease diagnostics and swine traits (Pathan et al., 2022; Piles et al., 2021). We believe that in biological systems, where small gene interactions can significantly interfere with the traits under evaluation (Meuwissen et al., 2016; Francisco et al., 2021), such factors must be assessed with care. Several authors using feature selection have reported promising results for genomic selection in plant and animal species (Aono et al., 2020; 2022; Li et al., 2022); however, the use of these reduced sets in new datasets, such as a new population or a test set, remains scarce. In practice, working with more stable predictors regardless of the MAF allows the breeder to have a more stable prediction accuracy in a range of scenarios, such as for traits controlled by a small number of QTLs or even when small populations are used as training sets, which can be obtained from random sampling, optimization, and minimizing the drift effect.

In contrast to the ML models, the traditional GS model used in this study (gBLUP) showed significant differences in predictive accuracy across all the traits due to the reduction in the number of SNPs by the MAF. As observed, all the traits showed decreases in PA when the dataset with an MAF of 0.20 was used, except for CW and RFM, whose PA values were intermediate in this dataset and higher than those at an MAF of 0.05 but lower than those at 0.10. With respect to the highest predictive accuracies with the traditional GS model, no significant differences were observed for TSS and CL, with MAFs of 0.05 and 0.10 showing significantly higher accuracies than the MAF of 0.20. Interestingly, for BL, CW, and RFM, the dataset with an MAF of 0.10 yielded substantially higher predictive accuracies than those obtained with the largest dataset (MAF of 0.05). Similar results were reported by Zhang et al. (2019), who reported that across different traditional GS models, such as additive gBLUP, additive-dominant gBLUP, and BayesR, a random reduction in the SNP set negatively affects predictive accuracy. According to Zhang et al. (2019), large sets of markers can also carry genotyping errors that negatively influence model estimates. In contrast, ML methods handle feature reduction well, enabling simpler models that often increase prediction accuracy (Aono et al., 2020; Phatan et al., 2022

### Implications for Plant Breeding

Over the past two decades, GS has been established as a robust alternative for optimizing breeding programs, resulting in significant increases in genetic gain for both plants and animals (Alemu et al., 2024). However, despite the advances in prediction models, the application of ML algorithms remains a fertile field of investigation, especially regarding the definition of optimized practices for SNP selection (Nguyen et al., 2015; Li et al., 2022). This study contributes to the debate by demonstrating that although traditional GS models achieve competitive PA, their performance is significantly affected by SNP set composition. In contrast, we observed that ML-based models exhibit greater stability under reduced marker density, maintaining consistent performance across external validation sets.

Beyond methodological advancements, this study presents direct implications for the productive sector by providing robust tools for grapevine breeding in Brazil. The implementation of early selection, grounded in our models, allows for a drastic reduction in operational costs and the interval between selection cycles, accelerating the release of superior cultivars (Jannink, 2010; Peixoto et al., 2024; Escamilla et al., 2025). The use of the IAC Germplasm Bank (BAG), a national reference center for *Vitis* spp., ensures that the trained models are representative of current elite genotype populations and of the selection populations of ongoing breeding programs. Additionally, we demonstrate that applying a simplified set of markers (MAF > 0.10) not only preserves accuracy but also optimizes the predictive performance for traits with complex genetic architectures, both chemical and morphological, making the use of GS more accessible and economically viable for national programs. Despite the demonstrated efficiency of the GS models presented here, their long-term sustainability remains to be validated. Future studies are needed to understand the necessity of recalibrating predictive models when reduced marker sets are used, ensuring that this reduction does not compromise the continuity of genetic gain in subsequent cycles (Jannink, 2010).

## Conclusion

Our results demonstrate a significant influence of MAF on traditional GS models; however, the same trend was not observed for the ML models. Overall, our findings suggest that an MAF of 0.10 is the most suitable threshold, as it frequently yields predictive accuracies superior to or equivalent to those of the other tested levels without compromising the trained model. With respect to the predictive models, SVR and ElasticNet achieved predictive accuracies that were significantly superior or equivalent to those of the others and remained unaffected by the reduction in MAF. The traditional GS model also showed high pr5edictive accuracy but proved highly sensitive to data reduction resulting from more restrictive MAF filtering. In summary, this work highlights the potential of GS in grapevine breeding programs, using both traditional and ML-based models, and emphasizes that the appropriate selection of molecular markers can determine the success of this tool.

## Author Contributions

F.R.F., G.L.O., G.F.N., R.F.-N., A.P.S., and M.F.M.F. conceptualized and designed the project. F.R.F. and R.F.-N. performed the data analyses. Data collection and experimental fieldwork were carried out by G.L.O., G.F.N., A.P.S., and M.F.M.F. The original draft writing and manuscript revision were structured and written by F.R.F., R.F.-N., and M.F.M.F. All authors reviewed and approved the final version of the manuscript.

## Acknowledgments

The authors would like to thank the Center for Plant Molecular Breeding (CeM²P) for the technical and structural support provided during most of this research.

## List of Abbreviations

AGB: Active Germplasm Bank
RFM: Rachis Fresh Mass
BL: Berry Length
BLUE: Best Linear Unbiased Estimate
BLUP: Best Linear Unbiased Prediction
CL: Cluster Length
CW: Cluster Width
EGGc: Expected Genetic Gain - Conventional Breeding
EGGpg: Expected Genetic Gain - Genomic Prediction
GBS: Genotyping-by-Sequencing
gBLUP: Genomic Best Linear Unbiased Prediction
GP: Genomic Prediction
GS: Genomic Selection
GWAS: Genome-Wide Association Studies
H²: Broad-Sense Heritability
IAC: Instituto Agronômico de Campinas
kNN: k-Nearest Neighbor
MAF: Minor Allele Frequency
MAS: Marker-Assisted Selection
MI: Maturation Index
ML: Machine Learning
NGS: Next-Generation Sequencing
PA: Predictive Accuracy
PCA: Principal Component Analysis
QTL: Quantitative Trait Locus
RFM: Rachis Fresh Mass
SNP: Single-Nucleotide Polymorphism
SVM: Support Vector Machine
SVR: Support Vector Regression
TSS: Total Soluble Solids
WGS: Whole-Genome Sequencing

