## Supplementary Material for "Assessing the Role of Marker Density and Minor Allele Frequency on Machine Learning–Driven Genomic Selection Accuracy in Grapevine"

### Supplementary Materials

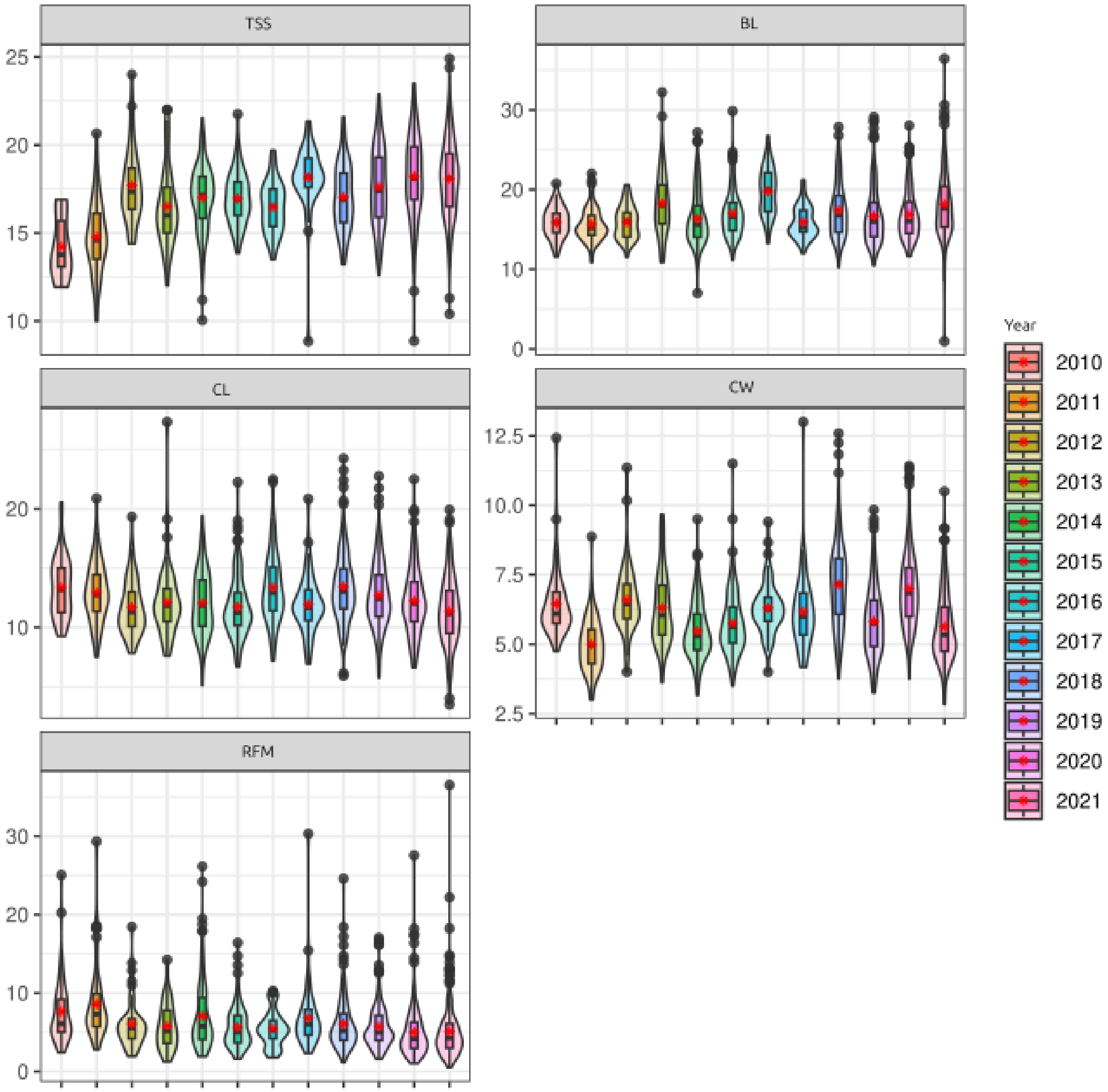


Supplementary Figure 1: Distribution of raw phenotypic data.

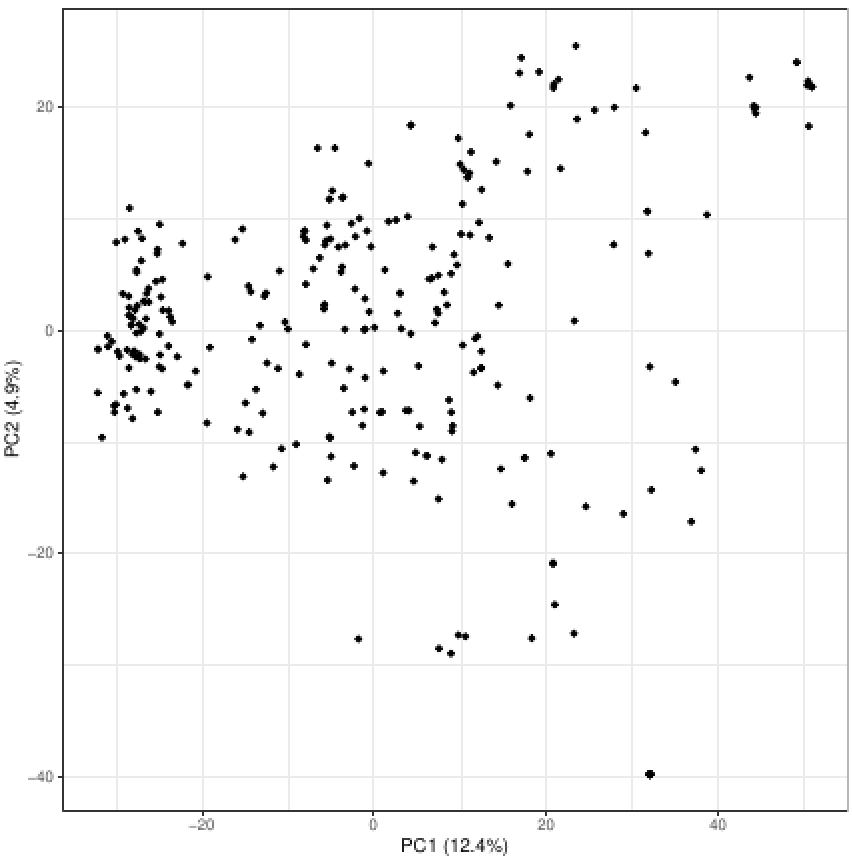


Supplementary Figure 2: Principal component analysis performed with the SNPs identified with MAF > 0.05.

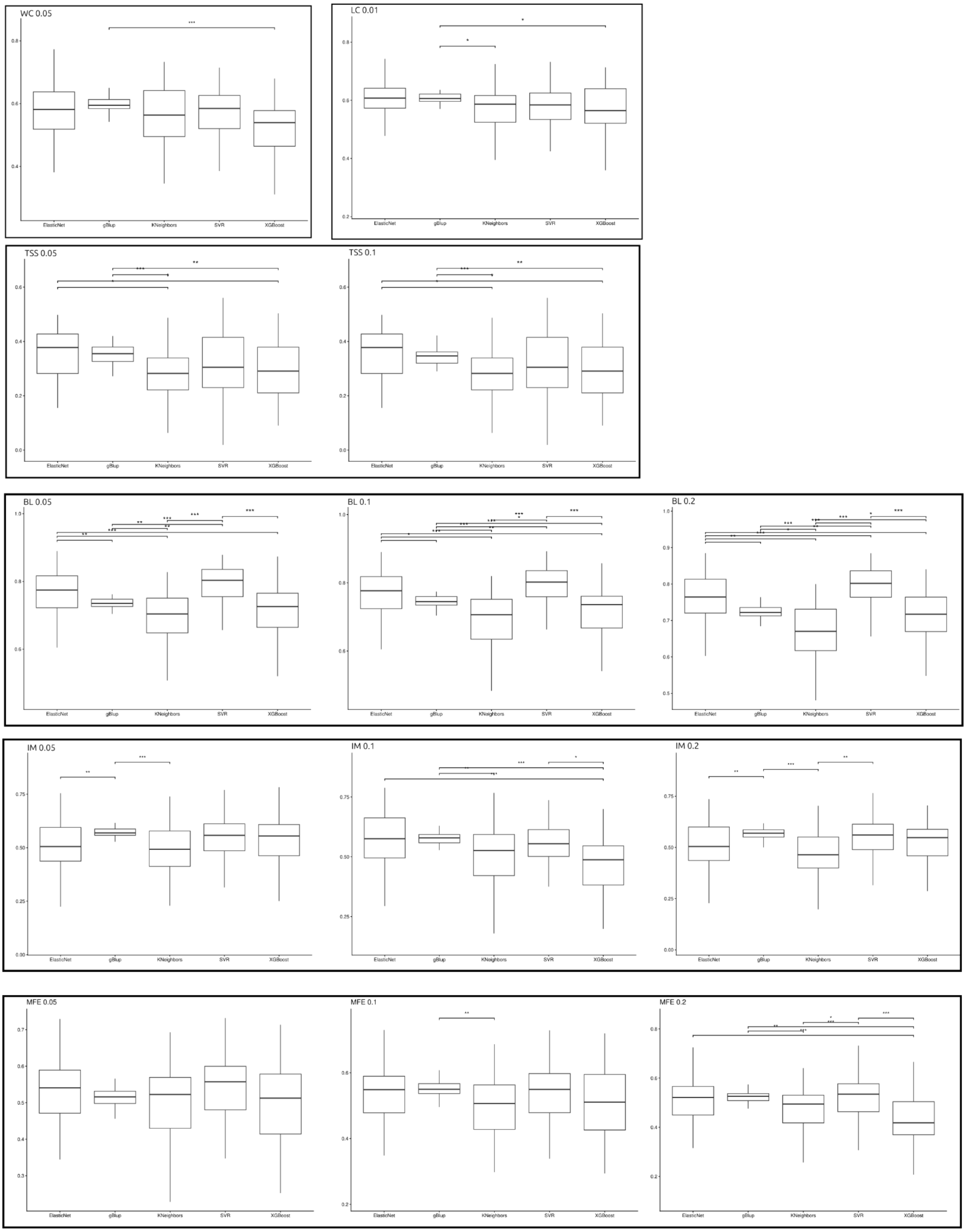


Supplementary Figure 3: Pairwise significance tests between models.

**Supplementary Table 1:** BLUEs of the traits evaluated for each individual

| id | TSS | CL | WC | BL | RFM | MI |
|---|---|---|---|---|---|---|
| 3 | -2.031 | -0.509 | 1.338 | 3.527 | 3.331 | -3.025 |
| 4 | -1.754 | 3.080 | 2.761 | 0.983 | 6.743 | 4.645 |
| 5 | 2.106 | -2.878 | 0.081 | 1.421 | -2.112 | 24.996 |
| 6 | 2.122 | 1.280 | 1.972 | 1.401 | 7.688 | 5.782 |
| 7 | 1.456 | -4.678 | -0.472 | 2.273 | -2.838 | 37.462 |
| 8 | 0.362 | -0.712 | -0.855 | -1.757 | 0.064 | -3.912 |
| 9 | -1.394 | 0.169 | 0.584 | 1.570 | 0.293 | -12.065 |
| 10 | 2.826 | -1.478 | -1.614 | -7.447 | -3.273 | 1.659 |
| 12 | -3.261 | -2.651 | -1.178 | -4.939 | -1.624 | -7.254 |
| 13 | -1.858 | -1.996 | -0.705 | -2.229 | -2.369 | -2.536 |
| 14 | -3.144 | -4.345 | -1.472 | 0.723 | -1.247 | -7.579 |
| 15 | -3.644 | 0.655 | -1.555 | -7.127 | 9.078 | -17.679 |
| 16 | -1.361 | -0.084 | 0.139 | -4.085 | 2.232 | -6.947 |
| 17 | -0.833 | -2.109 | -0.903 | -5.116 | 1.206 | -0.212 |
| 18 | -2.319 | -2.824 | -0.660 | -3.248 | -1.427 | -11.628 |
| 19 | -0.804 | -1.062 | -0.922 | -7.002 | 1.340 | -3.997 |
| 20 | 2.506 | -4.491 | -1.743 | -6.110 | -0.816 | 1.442 |
| 21 | -1.744 | -0.595 | 0.445 | 3.823 | -1.483 | 18.267 |
| 22 | -1.761 | 0.974 | 1.028 | -2.557 | 4.137 | -5.357 |
| 23 | -1.601 | -0.018 | -0.662 | -2.050 | -0.167 | -3.906 |
| 24 | -4.644 | -1.928 | 0.028 | -5.660 | 4.957 | -7.479 |
| 25 | -0.444 | -0.616 | 1.736 | 3.352 | 3.256 | 4.142 |
| 26 | 3.097 | 1.843 | -0.618 | -7.256 | 3.782 | 0.858 |
| 27 | -1.074 | -1.848 | -1.479 | -5.049 | -0.120 | -7.111 |
| 28 | 0.849 | 0.705 | 0.711 | -5.747 | 0.817 | 0.388 |
| 29 | 2.206 | 1.811 | 0.872 | 3.302 | 0.941 | 10.369 |
| 30 | -1.835 | 0.672 | 0.259 | 0.905 | 0.946 | -2.387 |
| 31 | -0.509 | -1.631 | 0.403 | 0.280 | 0.594 | 31.419 |
| 32 | -0.154 | -2.614 | -0.376 | -0.808 | -0.913 | 31.986 |
| 33 | -0.432 | -2.387 | 0.215 | -0.048 | -1.125 | 31.463 |
| 34 | -0.709 | -5.047 | -1.786 | -4.867 | -3.491 | 25.906 |
| 35 | -1.414 | -2.675 | 0.258 | 1.354 | 0.133 | 13.273 |
| 36 | -1.332 | -3.166 | -0.722 | -3.506 | -1.299 | -1.114 |
| 37 | -0.924 | -1.687 | 0.486 | 1.155 | -0.814 | 27.395 |
| 38 | -2.182 | -3.167 | -0.634 | -2.104 | -2.589 | 20.733 |
| 39 | -1.344 | -3.408 | -0.312 | 1.286 | -1.344 | 19.264 |
| 40 | -0.444 | -4.039 | -0.382 | 2.078 | -2.651 | 31.055 |
| 41 | -0.533 | -2.928 | -0.083 | 1.420 | -0.249 | 10.349 |
| 42 | 0.675 | 3.786 | -0.948 | -3.603 | 0.434 | 3.642 |
| 43 | -1.178 | -3.928 | -1.639 | -4.782 | -2.246 | 8.010 |
| 44 | -2.284 | -2.662 | -0.689 | -3.367 | -1.520 | -3.632 |
| 45 | -3.144 | -3.712 | -0.939 | -0.882 | -1.436 | 9.937 |
| 46 | -2.511 | -1.012 | 0.215 | -0.881 | 0.893 | 33.322 |
| 47 | 1.013 | -5.762 | -0.758 | -0.097 | -2.924 | 16.329 |
| 48 | 2.961 | -2.553 | -1.125 | -4.135 | -1.296 | 19.988 |

| | | | | | | |
|---|---|---|---|---|---|---|
| 49 | -0.053 | -4.769 | -0.562 | -0.333 | -2.589 | 2.593 |
| 50 | -2.230 | -3.536 | -1.472 | -3.427 | -3.653 | 8.712 |
| 51 | -1.994 | -4.414 | -1.333 | -2.181 | -3.332 | -0.696 |
| 52 | -2.933 | -1.262 | 1.070 | 0.873 | -0.044 | 11.697 |
| 53 | 0.584 | -2.172 | -1.555 | -4.273 | -2.608 | 7.502 |
| 54 | -2.053 | 0.724 | -0.235 | -3.649 | -0.297 | -3.665 |
| 56 | -0.082 | 1.665 | -0.597 | -3.042 | 0.828 | -3.157 |
| 57 | -1.003 | -2.845 | -0.535 | -2.373 | -0.838 | 9.131 |
| 58 | -0.144 | 0.271 | 0.745 | 0.878 | 0.186 | 3.529 |
| 59 | 1.256 | 0.905 | -0.422 | -1.987 | -0.741 | -1.723 |
| 60 | -0.044 | 2.005 | -0.622 | -1.970 | -1.422 | 10.539 |
| 61 | -2.011 | -0.320 | 1.357 | 2.988 | 4.174 | 8.871 |
| 62 | 1.618 | 0.509 | -0.889 | -7.744 | 5.580 | -9.307 |
| 63 | 0.056 | -1.178 | -0.448 | 2.528 | -0.593 | 4.972 |
| 64 | 1.982 | -0.529 | -0.455 | -2.676 | -1.715 | 1.259 |
| 65 | -0.644 | -2.245 | -0.889 | -3.177 | -2.853 | 7.176 |
| 66 | -0.794 | -0.062 | -0.389 | -5.897 | -3.517 | 1.943 |
| 67 | 1.213 | -2.288 | -0.210 | -4.646 | -1.710 | -2.504 |
| 68 | 1.117 | -2.859 | -1.139 | -4.357 | -2.218 | 8.498 |
| 69 | 0.117 | -3.365 | -0.028 | 0.651 | 0.655 | 2.308 |
| 70 | 1.322 | -1.035 | -0.829 | -2.379 | -2.071 | -5.273 |
| 71 | -0.319 | 1.030 | 1.632 | 0.806 | -0.061 | 5.005 |
| 72 | 0.713 | 3.631 | -0.805 | -3.437 | 1.528 | -0.342 |
| 73 | 0.056 | 3.821 | 0.715 | 7.213 | 4.274 | 6.892 |
| 74 | -0.500 | -3.928 | -1.833 | -3.394 | -3.619 | 6.153 |
| 75 | 0.531 | 0.759 | 0.153 | -8.140 | 4.037 | -16.290 |
| 76 | 1.346 | -3.862 | -1.239 | -5.584 | -2.005 | -14.898 |
| 77 | 1.722 | 2.488 | 2.139 | -5.594 | 3.462 | -10.994 |
| 79 | -1.211 | -1.095 | 0.472 | 0.456 | 0.778 | -2.445 |
| 80 | -1.028 | -2.199 | -0.597 | 2.235 | -1.561 | -8.178 |
| 81 | -0.644 | -0.699 | -0.139 | 0.306 | -2.549 | -3.120 |
| 82 | -2.897 | 0.904 | 0.104 | -1.783 | 0.938 | 5.824 |
| 83 | -1.155 | -2.746 | -0.133 | 2.923 | -3.063 | -4.656 |
| 84 | 0.956 | 2.022 | 0.195 | -4.660 | 4.370 | -10.198 |
| 85 | 0.376 | -0.095 | -0.672 | -4.078 | -2.559 | -2.588 |
| 86 | 0.556 | -0.212 | 0.228 | -3.917 | -0.438 | 5.722 |
| 87 | 0.456 | 3.966 | 3.667 | -3.456 | 8.316 | -12.061 |
| 88 | -2.578 | -2.137 | 0.486 | 3.009 | 0.487 | 1.045 |
| 89 | 1.836 | -0.588 | -0.172 | -6.046 | 0.457 | 1.790 |
| 90 | -2.684 | -0.978 | 0.845 | 8.665 | 0.521 | -5.610 |
| 91 | -0.659 | -0.059 | 0.314 | -0.503 | -0.428 | 23.519 |
| 92 | -2.266 | 0.627 | -0.611 | -3.471 | -0.904 | -5.341 |
| 93 | 1.847 | -0.442 | -1.319 | -9.983 | -2.070 | 1.745 |
| 94 | 0.326 | 5.422 | 0.668 | -7.469 | 6.848 | -8.750 |
| 95 | 1.681 | -1.949 | -0.930 | -6.202 | -1.574 | 0.270 |
| 96 | 0.956 | 0.183 | -0.368 | -7.024 | -0.663 | -4.512 |
| 97 | 0.926 | -2.712 | -1.772 | -7.614 | -1.855 | -8.509 |
| 98 | 0.681 | 0.842 | 1.132 | -5.506 | 1.979 | -11.066 |
| 99 | -0.136 | -1.817 | -1.597 | -4.916 | -1.413 | -2.812 |

| | | | | | | |
|---|---|---|---|---|---|---|
| 100 | -0.439 | 0.088 | 1.150 | -2.763 | 1.728 | -1.604 |
| 101 | -1.033 | -0.248 | 0.986 | -4.644 | 2.455 | -6.604 |
| 102 | -0.019 | -0.312 | 0.080 | -2.987 | 1.818 | -1.054 |
| 103 | 1.766 | 0.622 | -0.189 | -6.820 | -0.487 | -4.195 |
| 104 | 0.006 | 0.147 | -0.797 | -7.129 | -2.353 | 1.320 |
| 105 | -0.736 | -0.324 | -1.410 | -6.173 | -2.019 | -4.801 |
| 106 | 0.343 | -2.487 | -0.939 | -1.066 | -2.289 | 4.645 |
| 107 | -1.394 | 0.238 | -1.055 | -5.485 | 5.433 | -9.371 |
| 108 | 1.699 | 0.172 | -1.172 | -5.697 | -0.493 | 3.340 |
| 109 | 0.272 | -3.012 | -1.014 | -4.849 | -2.303 | -3.269 |
| 110 | -1.066 | -1.784 | -1.258 | -2.588 | -1.489 | 18.746 |
| 111 | 1.206 | 2.919 | -0.194 | -7.483 | 0.808 | -5.996 |
| 112 | -0.651 | 0.022 | 0.161 | -4.410 | 1.627 | -5.823 |
| 113 | 0.376 | -3.078 | 0.211 | -4.777 | -1.366 | 0.460 |
| 114 | 0.314 | -0.566 | -0.760 | -4.931 | -1.628 | 0.549 |
| 115 | -3.169 | 6.645 | 1.327 | 2.494 | 2.011 | -6.497 |
| 116 | -1.571 | -1.820 | 0.678 | 1.771 | 0.364 | -4.140 |
| 117 | 2.409 | -1.480 | 0.232 | -5.061 | 2.780 | 13.904 |
| 118 | 1.576 | -1.795 | -0.055 | -6.980 | -0.393 | -5.550 |
| 119 | 2.506 | -3.595 | -3.555 | -10.194 | -3.283 | -8.129 |
| 120 | 0.396 | -2.545 | -2.705 | -7.974 | -2.853 | 10.032 |
| 121 | 0.251 | -3.678 | -2.240 | -7.209 | -1.265 | 3.936 |
| 122 | 0.961 | 0.363 | -1.153 | -5.844 | -0.112 | 4.903 |
| 123 | 0.476 | -3.103 | -0.972 | -8.870 | -2.710 | -0.120 |
| 124 | 1.089 | -0.870 | -1.314 | -8.227 | -1.301 | 2.033 |
| 125 | -5.944 | 2.072 | -1.472 | -9.277 | -3.742 | -15.128 |
| 126 | 0.756 | -0.637 | -0.111 | -7.008 | -1.899 | -5.913 |
| 128 | 2.696 | 2.372 | 0.728 | -6.537 | 2.309 | 2.490 |
| 129 | 2.282 | 2.338 | 0.661 | -6.294 | 2.854 | -4.023 |
| 131 | -1.400 | -1.873 | -1.139 | -1.633 | -2.956 | 5.881 |
| 133 | -0.900 | 1.516 | 0.667 | -3.227 | 7.863 | 1.537 |
| 134 | 2.234 | -1.052 | -0.307 | -2.247 | -1.010 | 10.467 |
| 135 | -0.697 | 1.238 | 0.382 | -4.163 | 1.173 | -2.726 |
| 136 | -2.061 | -0.481 | 1.842 | 5.626 | 0.517 | -9.485 |
| 137 | -0.244 | -4.928 | -2.622 | -2.887 | -3.810 | 5.270 |
| 138 | -3.619 | -0.541 | -0.360 | 3.240 | 3.619 | -4.654 |
| 139 | -0.124 | -1.512 | 0.911 | -1.827 | -0.617 | 1.337 |
| 140 | 0.856 | -3.262 | -0.639 | 0.990 | -3.382 | -4.404 |
| 141 | -0.544 | 2.572 | -0.889 | 0.206 | 3.745 | -5.428 |
| 142 | -0.819 | 2.737 | -0.235 | -2.624 | 2.212 | -11.479 |
| 143 | -1.834 | -1.345 | 0.245 | 0.775 | 1.304 | -7.730 |
| 144 | -3.094 | -1.658 | -0.335 | 2.955 | 2.539 | -5.830 |
| 145 | -0.984 | -2.053 | -0.247 | 1.923 | -0.033 | 9.099 |
| 146 | -3.530 | -4.547 | -1.186 | -0.701 | -3.515 | -10.337 |
| 148 | -0.478 | -2.666 | -0.523 | -3.749 | -2.828 | 15.236 |
| 150 | -0.744 | -3.428 | -1.005 | -6.804 | -1.996 | -10.413 |
| 151 | 1.486 | -2.912 | -1.545 | -5.154 | -3.309 | 8.590 |
| 152 | 0.856 | -2.449 | -0.555 | -4.490 | -0.203 | -1.326 |
| 153 | -4.144 | 3.947 | 0.403 | 4.973 | 13.734 | -12.229 |

| | | | | | | |
|---|---|---|---|---|---|---|
| 156 | -1.419 | -2.901 | -0.153 | 0.319 | -0.681 | 8.745 |
| 157 | 3.989 | -2.345 | -0.666 | -8.158 | -0.744 | -8.529 |
| 158 | 0.156 | -1.553 | -1.910 | -4.848 | 0.451 | -11.178 |
| 159 | -1.216 | 0.209 | -0.651 | -1.768 | -2.138 | -6.315 |
| 160 | 2.306 | -3.928 | -2.055 | -6.052 | 4.932 | -3.204 |
| 161 | -0.764 | -1.895 | 0.095 | -1.160 | -0.946 | -3.750 |
| 162 | -2.269 | -1.414 | -0.139 | 1.623 | -2.875 | -3.038 |
| 163 | -0.019 | -4.573 | -3.220 | -9.996 | -3.521 | -7.932 |
| 166 | -1.569 | -6.720 | -1.930 | -5.248 | -4.021 | -16.352 |
| 167 | -2.044 | -2.928 | 0.361 | -0.510 | -2.598 | -15.277 |
| 170 | -4.444 | -7.262 | -2.639 | -6.110 | -4.374 | -19.877 |
| 174 | -3.744 | -6.512 | -0.389 | -2.127 | -4.342 | -0.429 |
| 176 | -7.244 | -6.528 | -0.722 | -6.507 | -4.710 | -15.427 |
| 177 | 0.456 | -8.178 | -2.805 | -2.294 | -5.080 | 22.018 |
| 178 | -3.544 | -7.428 | -0.722 | -6.694 | -5.267 | 20.669 |
| 181 | 1.156 | -9.928 | -3.889 | -12.527 | -4.818 | -22.326 |
| 183 | -1.471 | -3.465 | -1.163 | -1.876 | -1.754 | 10.276 |
| 184 | -0.701 | -4.268 | -1.984 | -5.067 | -3.426 | 15.333 |
| 185 | -0.502 | -4.565 | -1.340 | -3.703 | -3.339 | 23.802 |
| 186 | -1.173 | -0.935 | -1.030 | -3.880 | -2.317 | 2.258 |
| 187 | 1.847 | -1.112 | -0.583 | -5.486 | 0.342 | 0.547 |
| 188 | 0.625 | -1.090 | -0.764 | -1.528 | -2.545 | -1.696 |
| 189 | 0.518 | -2.087 | -2.271 | -8.266 | -0.437 | -12.943 |
| 190 | -2.096 | -3.053 | -2.186 | -7.308 | -2.144 | -7.114 |
| 191 | 0.881 | -2.928 | -0.778 | -3.031 | -1.559 | -6.772 |
| 192 | 0.660 | -0.488 | -1.185 | -3.550 | -1.201 | 2.554 |
| 193 | -1.368 | -3.613 | -1.504 | -8.035 | 3.197 | -10.826 |
| 194 | 0.980 | -3.702 | -1.573 | -7.864 | -1.255 | -4.413 |
| 195 | -2.594 | -1.943 | -0.530 | -0.980 | -1.311 | 5.407 |
| 196 | -1.016 | -2.921 | -1.135 | -7.217 | 1.994 | -10.993 |
| 197 | -0.601 | 1.530 | -0.657 | -5.892 | 3.818 | -2.540 |
| 198 | 1.459 | -0.926 | -0.489 | -7.077 | 1.248 | -7.772 |
| 199 | 0.562 | -2.538 | -0.717 | -6.053 | 0.310 | -2.432 |
| 200 | 2.229 | 2.446 | 0.507 | -6.674 | 4.956 | -4.178 |
| 201 | 1.515 | -1.910 | -0.067 | -3.910 | 1.442 | 8.308 |
| 202 | -0.961 | -3.200 | -1.154 | -6.831 | -1.230 | -6.349 |
| 203 | -1.821 | -4.357 | -1.121 | -0.645 | -2.730 | 39.265 |
| 204 | 0.413 | -2.672 | -0.704 | -3.752 | -1.837 | 20.182 |
| 205 | 3.310 | -1.100 | -0.679 | -4.649 | 0.676 | 0.683 |
| 206 | 0.801 | -2.732 | -0.811 | -7.250 | 0.685 | -3.090 |
| 207 | -0.573 | -1.321 | -0.977 | -4.792 | -1.236 | 0.897 |
| 208 | -1.709 | -1.736 | -0.204 | -5.949 | 0.072 | -8.145 |
| 209 | 2.181 | -3.783 | -1.589 | -7.072 | -0.649 | 5.404 |
| 210 | 0.654 | -2.996 | -0.805 | -7.281 | -0.238 | 1.723 |
| 211 | 0.291 | 1.703 | -0.746 | -3.169 | -1.264 | -1.556 |
| 212 | 1.681 | 0.697 | 0.625 | -3.606 | -0.050 | 28.262 |
| 213 | 4.563 | -0.607 | -1.436 | -6.477 | -0.991 | 1.774 |
| 214 | 0.898 | -3.466 | -0.842 | -4.911 | -2.217 | -2.849 |
| 215 | 0.188 | -3.749 | -0.676 | -4.719 | -1.338 | -9.437 |

| | | | | | | |
|---|---|---|---|---|---|---|
| 216 | 2.506 | -0.614 | -0.257 | -6.370 | 1.646 | 3.303 |
| 217 | 0.286 | 2.666 | 1.740 | 5.573 | 5.064 | 13.637 |
| 218 | -1.247 | -2.393 | -1.085 | -5.852 | -0.743 | -4.964 |
| 219 | -1.287 | -2.881 | -1.058 | -6.366 | -0.511 | -9.886 |
| 220 | -0.186 | -3.477 | -0.553 | -6.385 | -1.450 | -3.334 |
| 221 | -0.179 | -2.928 | -1.135 | -6.623 | -1.336 | -3.117 |
| 222 | -0.353 | -2.386 | -0.410 | -3.827 | 1.904 | -7.692 |
| 223 | -0.544 | -2.185 | -0.997 | -5.444 | 3.527 | -6.427 |
| 224 | 0.681 | -4.356 | -0.694 | -4.123 | -0.352 | -3.122 |
| 225 | -0.562 | -3.166 | -0.455 | -6.161 | 3.402 | 0.465 |
| 226 | 0.856 | -4.001 | -1.014 | -6.592 | 0.639 | 7.193 |
| 227 | -0.069 | -5.037 | -1.604 | -6.384 | -1.620 | -6.804 |
| 228 | -1.119 | -2.046 | -0.772 | -5.863 | 1.669 | -7.319 |
| 229 | -0.825 | -2.928 | -1.060 | -5.028 | 1.624 | -8.977 |
| 230 | 2.012 | -3.079 | 0.204 | -6.827 | 5.668 | -4.345 |
| 231 | 1.477 | -7.847 | -2.519 | -7.365 | -3.024 | -1.562 |
| 232 | 0.806 | -4.172 | -0.847 | -3.345 | -0.694 | 5.268 |
| 233 | -0.161 | -6.039 | -1.535 | -6.545 | -1.193 | -1.294 |
| 234 | 0.441 | -4.247 | -0.421 | -4.337 | 2.317 | -6.758 |
| 235 | 0.646 | -6.597 | -2.175 | -6.803 | -2.761 | -1.226 |
| 236 | 0.356 | -0.459 | -0.272 | -2.261 | 3.294 | -2.370 |
| 237 | 2.674 | -6.203 | -2.242 | -5.224 | -1.708 | -0.249 |
| 238 | -1.995 | 4.665 | 2.701 | -4.272 | 19.713 | 2.562 |
| 240 | -1.051 | 1.305 | 0.761 | -2.865 | 5.204 | 2.777 |
| 241 | 0.456 | -3.206 | -1.500 | -5.199 | -1.942 | 4.852 |
| 242 | 2.889 | -6.275 | -1.041 | -3.863 | -3.443 | 7.786 |
| 243 | 1.089 | -7.373 | -2.000 | -6.488 | -2.915 | 3.920 |
| 244 | -0.744 | 1.322 | 1.403 | -3.827 | 3.394 | 1.345 |
| 245 | -4.178 | -0.873 | -1.764 | -6.485 | 0.280 | -4.047 |
| 246 | -2.811 | -5.539 | -1.472 | -4.316 | -2.389 | -8.610 |
| 247 | -4.094 | -2.553 | -0.597 | -7.044 | 3.024 | -5.553 |
| 248 | -0.794 | 1.843 | -0.201 | -4.923 | 2.025 | -0.553 |
| 249 | -2.894 | -4.137 | -1.097 | -7.002 | -0.326 | -10.602 |
| 250 | -1.570 | 0.217 | 0.026 | -4.786 | 2.094 | -8.406 |
| 252 | -4.144 | -3.262 | -0.111 | -1.088 | -0.563 | -6.561 |
| 253 | 1.256 | -6.762 | -2.889 | -8.077 | -3.138 | 9.521 |
| 254 | 1.989 | -1.678 | 0.500 | -3.105 | -1.016 | 1.988 |
| 255 | -0.144 | -5.275 | -1.416 | -7.891 | -2.233 | -6.861 |
| 256 | 0.602 | -4.174 | -1.002 | -3.986 | -0.380 | -12.757 |
| 258 | 0.456 | -2.789 | -0.278 | -4.160 | 0.004 | -2.344 |
| 259 | 1.106 | -2.895 | -0.672 | -3.465 | 1.255 | -3.154 |
| 260 | -0.644 | -1.595 | -2.505 | -7.540 | -0.885 | -3.878 |
| 261 | -1.244 | -7.262 | -1.555 | -6.794 | -3.972 | -7.177 |
| 262 | 0.389 | -2.206 | -0.472 | -3.016 | 1.225 | -4.044 |
| 263 | 0.556 | -8.678 | -2.722 | -20.077 | 8.841 | -4.427 |
| 264 | -1.411 | -4.978 | -0.844 | -2.296 | -0.347 | -0.312 |
| 266 | -1.244 | -3.595 | -1.305 | -4.777 | -0.812 | -1.695 |
| 267 | -0.744 | -3.206 | 0.278 | -2.821 | 1.335 | -2.795 |
| 268 | -0.044 | -3.289 | -0.444 | -3.110 | 0.074 | -4.095 |

| | | | | | | |
|---|---|---|---|---|---|---|
| 269 | -2.644 | -4.345 | -1.826 | -3.989 | -2.037 | -12.152 |
| 270 | 3.522 | -0.762 | 0.250 | -7.560 | 3.338 | 6.987 |
| 272 | 2.081 | -4.665 | -1.738 | -4.463 | -0.494 | 1.021 |
| 273 | 0.272 | 0.685 | 0.133 | -4.835 | 8.018 | -4.013 |
| 274 | -2.364 | -6.262 | -2.705 | -5.397 | -3.277 | -13.092 |
| 275 | -0.216 | -4.095 | -1.180 | -6.633 | 1.703 | -11.214 |
| 276 | -0.328 | -2.977 | -1.379 | -6.874 | 0.291 | -10.288 |
| 277 | 1.306 | -0.803 | -1.572 | -4.409 | 1.625 | 0.572 |
| 278 | -0.479 | -5.112 | -1.714 | -5.627 | -1.773 | -13.413 |
| 279 | 1.434 | -2.274 | -0.336 | -7.426 | 0.964 | -10.568 |
| 280 | 0.017 | -4.187 | -0.435 | -6.559 | 2.879 | -7.298 |
| 281 | 2.231 | -3.991 | -1.014 | -6.056 | -0.504 | 4.058 |
| 282 | 0.965 | -3.185 | -1.083 | -3.519 | -0.936 | 3.912 |
| 284 | 0.145 | -1.373 | -0.403 | -1.645 | -1.370 | 1.276 |
| 285 | 0.289 | -5.144 | -1.826 | -7.153 | -2.240 | -9.432 |
| 286 | 0.864 | -1.449 | -1.430 | -3.814 | -0.537 | 2.353 |
| 287 | 0.914 | -5.202 | -1.603 | -5.826 | -2.495 | 4.179 |
| 288 | -1.024 | -2.595 | -1.522 | -6.169 | -0.234 | -6.899 |
| 289 | -0.401 | -4.357 | -1.371 | -4.859 | -2.117 | -6.487 |
| 290 | 0.468 | -3.024 | -0.327 | -4.227 | -0.379 | -7.088 |
| 291 | -1.053 | -3.491 | -1.712 | -7.142 | -0.569 | -1.866 |
| 292 | -1.197 | -1.960 | -0.482 | -7.546 | 0.484 | -7.447 |
| 293 | -0.232 | 0.051 | -0.878 | -5.568 | -1.115 | -8.124 |
| 294 | -0.403 | -2.230 | -2.061 | -6.271 | -2.998 | -8.605 |
| 295 | 0.341 | -0.766 | -0.287 | -7.495 | 1.148 | -3.148 |
| 296 | 4.136 | -1.570 | -1.922 | -7.920 | 0.562 | -0.936 |
| 297 | 0.762 | -1.278 | -1.080 | -5.175 | 0.709 | -0.044 |
| 298 | -0.787 | -3.331 | -0.868 | -6.855 | -0.256 | -3.513 |
| 299 | 0.975 | -0.123 | -0.007 | -8.926 | 2.792 | -7.283 |
| 300 | -0.604 | -0.862 | -1.147 | -5.447 | -1.093 | -3.646 |
| 302 | 2.041 | 1.397 | -0.239 | -7.282 | 3.232 | -2.816 |
| 303 | 1.653 | -0.560 | 0.000 | -7.385 | 3.480 | -2.601 |
| 304 | -2.511 | 2.961 | 1.389 | -5.721 | 15.484 | -0.835 |
| 306 | 5.556 | -3.720 | -2.597 | -7.102 | -1.978 | 8.943 |
| 307 | 1.656 | -4.456 | -1.222 | -7.383 | -1.543 | -6.013 |
| 308 | -0.111 | 3.883 | 1.717 | 7.873 | 7.247 | 12.617 |
| 309 | -2.044 | 2.738 | 1.861 | 8.156 | 1.818 | 23.017 |
| 310 | -1.261 | 2.391 | -0.611 | -3.827 | 0.527 | 0.760 |
| 311 | -3.144 | -2.039 | -0.500 | -4.444 | 0.883 | -10.613 |

Supplementary Table 2: Hyperparameters tested in the machine learning models using the grid search method

| **Algorithm** | **Hyperparameters** | **Search Space** |
|---|---|---|
| ElasticNet | alpha, l1_ratio, max_iter | {0.1, 1, 10, 100}, {0.1, 0.5, 0.9}, {15000} |
| SVR | C, kernel, gamma | {0.1, 1, 10}, {linear, rbf}, {scale, auto} |
| Gradient Boosting | n_estimators, learning_rate, max_depth | {50, 100}, {0.001, 0.005, 0.01, 0.05, 0.1, 1.0}, {3, 5} |
| K-Neighbors | n_neighbors, weights | {3, 5, 7}, {uniform, distance} |
| XGBoost | n_estimators, learning_rate, max_depth | {5, 10, 20, 50, 100}, {0.001, 0.01, 0.05, 0.1}, {3, 5, 7, 10} |

| **Supplementary Table 3:** Mean predictive accuracies for the tested predictive models | | | | | | | |
|---|---|---|---|---|---|---|---|
| Model | MAF | BL | CL | CW | MI | RFM | TSS |
| ElasticNet | 0.05 | 0.770 | 0.600 | 0.560 | 0.530 | 0.520 | 0.340 |
| gBlup | 0.05 | 0.730 | 0.590 | 0.600 | 0.570 | 0.510 | 0.350 |
| KNeighbors | 0.05 | 0.700 | 0.540 | 0.550 | 0.510 | 0.480 | 0.290 |
| SVR | 0.05 | 0.790 | 0.590 | 0.580 | 0.550 | 0.540 | 0.300 |
| XGBoost | 0.05 | 0.720 | 0.570 | 0.540 | 0.490 | 0.500 | 0.300 |
| ElasticNet | 0.1 | 0.770 | 0.600 | 0.550 | 0.550 | 0.520 | 0.340 |
| gBlup | 0.1 | 0.750 | 0.610 | 0.590 | 0.580 | 0.550 | 0.340 |
| KNeighbors | 0.1 | 0.690 | 0.550 | 0.540 | 0.490 | 0.480 | 0.290 |
| SVR | 0.1 | 0.790 | 0.590 | 0.570 | 0.560 | 0.540 | 0.300 |
| XGBoost | 0.1 | 0.730 | 0.570 | 0.560 | 0.500 | 0.500 | 0.300 |
| ElasticNet | 0.2 | 0.760 | 0.600 | 0.550 | 0.530 | 0.500 | 0.310 |
| gBlup | 0.2 | 0.720 | 0.600 | 0.580 | 0.570 | 0.520 | 0.290 |
| KNeighbors | 0.2 | 0.670 | 0.550 | 0.540 | 0.480 | 0.450 | 0.280 |
| SVR | 0.2 | 0.790 | 0.590 | 0.570 | 0.550 | 0.520 | 0.310 |
| XGBoost | 0.2 | 0.720 | 0.580 | 0.560 | 0.500 | 0.410 | 0.320 |

| **Supplementary Table 4:** Overall significance test between the tested models. | | | | | | |
|---|---|---|---|---|---|---|
| MAF | BL | CL | CW | RFM | IM | TSS |
| 0.05 | 5.69E-14 | 0.001014 | 0.001014 | 0.03319 | 0.003608 | 0.0006189 |
| 0.1 | 9.94E-16 | 0.094350 | 0.020503 | 0.01166 | 3.61E-06 | 0.0015870 |
| 0.2 | 2.20E-16 | 0.238100 | 0.198300 | 1.10E-09 | 2.05E-05 | 0.1750009 |

| **Supplementary Table 5:** P values for predictive accuracy of each model across different MAF-based datasets | | | | | |
|---|---|---|---|---|---|
| Model | BL | CL | CW | RFM | TSS |
| ElasticNet | 0.896 | 0.994 | 0.766 | 0.440 | 0.255 |
| gBlup | 0.000 | 0.000 | 0.003 | 0.000 | 0.000 |
| Kneighbors | 0.191 | 0.889 | 0.854 | 0.307 | 0.892 |
| SVR | 0.982 | 0.988 | 0.953 | 0.590 | 0.993 |
| XGBoost | 0.974 | 0.686 | 0.723 | 0.000 | 0.377 |

| **Supplementary Table 6**: Predictive accuracy of the models on the test set | | | | | | | |
|---|---|---|---|---|---|---|---|
| Model | MAF | BL | CL | IM | RFM | WC | TSS |
| ElasticNet | 0.05 | 0.81 | 0.76 | 0.77 | 0.77 | 0.75 | 0.42 |
| gBlup | 0.05 | 0.82 | 0.66 | 0.70 | 0.75 | 0.59 | 0.30 |
| KNeighbors | 0.05 | 0.79 | 0.62 | 0.78 | 0.64 | 0.65 | 0.35 |
| SVR | 0.05 | 0.84 | 0.76 | 0.85 | 0.82 | 0.76 | 0.35 |

| | | | | | | | |
|---|---|---|---|---|---|---|---|
| XGBoost | 0.05 | 0.82 | 0.72 | 0.74 | 0.66 | 0.68 | 0.30 |
| ElasticNet | 0.1 | 0.82 | 0.75 | 0.78 | 0.77 | 0.74 | 0.38 |
| gBlup | 0.1 | 0.75 | 0.67 | 0.59 | 0.61 | 0.64 | 0.24 |
| KNeighbors | 0.1 | 0.77 | 0.60 | 0.78 | 0.63 | 0.66 | 0.39 |
| SVR | 0.1 | 0.84 | 0.76 | 0.86 | 0.82 | 0.76 | 0.37 |
| XGBoost | 0.1 | 0.80 | 0.70 | 0.71 | 0.74 | 0.73 | 0.53 |
| ElasticNet | 0.2 | 0.81 | 0.75 | 0.75 | 0.75 | 0.61 | 0.40 |
| gBlup | 0.2 | 0.84 | 0.58 | 0.67 | 0.61 | 0.67 | 0.72 |
| KNeighbors | 0.2 | 0.71 | 0.63 | 0.69 | 0.65 | 0.68 | 0.30 |
| SVR | 0.2 | 0.85 | 0.75 | 0.84 | 0.82 | 0.74 | 0.42 |
| XGBoost | 0.2 | 0.84 | 0.69 | 0.72 | 0.67 | 0.57 | 0.51 |
